## Supplementary material for "An interactive meta-analysis of MRI biomarkers of myelin": Interactive figures (Jupyter Notebook+HTML): interactive_figures_rev1.ipynb

```
# %% [code]
# Run this cell in Google Colab to install the required packages and retrieve the data
!wget https://raw.githubusercontent.com/matteomancini/myelin-meta-analysis/master/requirements.txt
!wget https://github.com/matteomancini/myelin-meta-analysis/raw/master/database.xlsx
!pip install -r requirements.txt
!echo "install.packages(\"metafor\", repos=\"https://cran.rstudio.com\")" | R --no-save
!echo "install.packages(\"multcomp\", repos=\"https://cran.rstudio.com\")" | R --no-save

# %% [code]
import numpy as np
import pandas as pd

import plotly.graph_objects as go
import plotly.express as px
import plotly.colors
from plotly.subplots import make_subplots

from rpy2.robjects.packages import importr
import rpy2.robjects

# %% [code]
screening_info = ['Records obtained from the Medline database',
                  'Records obtained from previous reviews',
                  """
                    Exclusion critera:<br>
                    - work relying only on MRI;<br>
                    - work relying only on histology or equivalent approach;<br>
                    - work reporting only qualitative comparisons.
                  """,
                  'Records selected for full-text evaluation',
                  """
                    Exclusion criteria:<br>
                    - studies using MRI-based measures in arbitrary units;<br>
                    - studies using measures of variation in myelin content;<br>
                    - studies using arbitrary assessment scales;<br>
                    - studies comparing absolute measures of myelin with relative measures;<br>
                    - studies reporting other quantitative measures than correlation or R^2 values;<br>
                    - studies comparing histology from one dataset and MRI from a different one.
                  """,
                  'Studies selected for literature overview',
                  """
                    Exclusion criteria:<br>
                     - not providing an indication of both number of subjects and number of ROIs.
                  """]

fig1 = go.Figure(data=[go.Sankey(
    arrangement = "freeform",
    node = dict(
      pad = 15,
      thickness = 20,
      line = dict(color = "black", width = 0.5),
      label = ["Main records identified (database searching)",
               "Additional records (reviews)",
               "Records screened",
               "Records excluded",
               "Full-text articles assessed for eligibility",
               "Full-text articles excluded",
               "Studied included in the literature overview",
               "Studies included in the meta-analysis"],
      x = [0, 0, 0.4, 0.6, 0.5, 0.8, 0.7, 1],
      y = [0, 0, 0.5, 0.8, 0.15, 0.05, 0.4, 0.6],
      hovertemplate = "%{label}<extra>%{value}</extra>",
      color = ["darkblue","darkblue","darkblue","darkred","darkgreen","darkred","darkgreen","darkgreen"]
    ),
    link = dict(
      source = [0, 1, 2, 2, 4, 4, 6],
      target = [2, 2, 3, 4, 5, 6, 7],
      value = [688, 1, 597, 92, 34, 58, 43],
      customdata = screening_info,
      hovertemplate = "%{customdata}",
  ))])

fig1.update_layout(title='Figure S1: Review methodology',
                   width=1000,
                   height=500,
                   font_size=12)
fig1.show()

# %% [code]
info = pd.read_excel('database.xlsx', sheet_name='Details')

year_str = info['Year'].astype(str)
info['Study'] = info['First author'] + ' et al., ' + year_str
info['Study'] = info.groupby('Study')['Study'].apply(lambda n: n+list(map(chr,np.arange(len(n))+97))
                                                     if len(n)>1 else n)
info['Number of studies'] = np.ones((len(info),1))
info = info.sort_values('Study')

info['Link'] = info['DOI']
info['Link'].replace('http',"""<a style='color:white' href='http""",
                    inplace=True, regex=True)
info['Link'] = info['Link'] + """'>->Go to the paper</a>"""

fields = ['Approach', 'Magnetic field', 'MRI measure(s)',
          'Histology/microscopy measure', 'Specific structure(s)']
info['Summary'] = info['Link'] + '<br><br>'
for i in fields:
    info['Summary'] = info['Summary'] + i + ': ' + info[i].astype(str) + '<br><br>'

args = dict(data_frame=info, values='Number of studies',
            color='Number of studies', hover_data='',
            path=['Focus', 'Tissue condition', 'Human/animal', 'Condition', 'Study'],
            color_continuous_scale='Viridis')
args = px._core.build_dataframe(args, go.Treemap)
treemap_df = px._core.process_dataframe_hierarchy(args)['data_frame']

fig2 = go.Figure(go.Treemap(
        ids=treemap_df['id'].tolist(),
        labels=treemap_df['labels'].tolist(),
        parents=treemap_df['parent'].tolist(),
        values=treemap_df['Number of studies'].tolist(),
        branchvalues='total',
        text=info['Summary'],
        hoverinfo='label',
        textfont=dict(
            size=15,
        )
    )
)

fig2 = fig2.update_layout(
    autosize=False,
    width=900,
    height=600,
    margin=dict(
        l=0,
        r=0,
        b=30,
        t=60,
    )
)

fig2.update_layout(
    title='Figure S2: Literature survey overview')

fig2.show()

# %% [code]
df = pd.DataFrame()
data = pd.read_excel('database.xlsx', sheet_name='R^2')

measures = data.columns[1:]
for _, row in data.iterrows():
    measure_avail = {m:value for m, value in zip(measures, row.tolist()[1:])
                    if not np.isnan(value)}
    for m in measure_avail.keys():
        df = df.append([[row.DOI, m, measure_avail[m],
                         *info[info.DOI==row.DOI].values.tolist()[0][1:]]])
df.columns = ['DOI', 'Measure', 'R^2', *info.columns[1:]]

df['ROI per subject'] = pd.to_numeric(df['ROI per subject'], errors='coerce')
df['Subjects'] = pd.to_numeric(df['Subjects'], errors='coerce')
df = df.dropna(subset=['ROI per subject', 'Subjects'])
df = df[df['ROI per subject']<100]
df['Sample points'] = df['ROI per subject'] * df['Subjects']

df=df.sort_values(by=['Measure'])

filtered_df=df[df.Focus=='Brain'].copy()

measure_type = {'Diffusion':['RD', 'AD', 'FA', 'MD',
                'AWF', 'RK', 'RDe', 'MK'],
                'Magnetization transfer':['MTR',
                'ihMTR', 'MTR-UTE', 'MPF', 'MVF-MT',
                'R1f', 'T2m', 'T2f', 'k_mf','k_fm'],
                'T1 relaxometry':['T1'], 'T2 relaxometry':['T2', 'MWF', 'MVF-T2'],
                'Other':['QSM', 'R2*', 'rSPF', 'MTV',
                'T1p', 'T2p', 'RAFF', 'PD', 'T1sat']}

color_dict = {m:plotly.colors.qualitative.Bold[n]
              for n,m in enumerate(measure_type.keys())}

hover_text = []
bubble_size = []

for index, row in filtered_df.iterrows():
    hover_text.append(('Measure: {measure}<br>'+
                      'Number of subjects: {subjects}<br>'+
                      'ROIs per subject: {rois}<br>'+
                      'Total number of samples: {samples}').format(measure=row['Measure'],
                                            subjects=row['Subjects'],
                                            rois=row['ROI per subject'],
                                            samples=row['Sample points']))
    bubble_size.append(2*np.sqrt(row['Sample points']))

filtered_df['Details'] = hover_text
filtered_df['Size'] = bubble_size

fig3 = go.Figure()

for m in measure_type.keys():
    df_m = filtered_df[filtered_df['Measure'].isin(measure_type[m])]
    fig3.add_trace(go.Scatter(
                    x=df_m['Measure'],
                    y=df_m['R^2'],
                    text='Study: ' + 
                        df_m['Study']+ '<br>' + df_m['Details'],
                    mode='markers',
                    line = dict(color = 'rgba(0,0,0,0)'),
                    marker = dict(color=color_dict[m]),
                    marker_size = df_m['Size'],
                    opacity=0.6,
                    name=m
                    ))
    
fig3.update_layout(
    title='Figure S3: R<sup>2</sup> between MRI and histology across measures',
    xaxis=dict(title='MRI measure'),
    yaxis=dict(title='R<sup>2</sup>'),
    autosize=False,
    width=900,
    height=600
)

fig3.show()

# %% [code]
filtered_df=filtered_df.sort_values(by=['Study', 'Measure'])

args = dict(data_frame=filtered_df, values='Sample points',
            color='R^2', hover_data='',
            path=['Measure', 'Study'],
            color_continuous_scale='Viridis')
args = px._core.build_dataframe(args, go.Treemap)
treemap_df = px._core.process_dataframe_hierarchy(args)['data_frame']

fig4 = go.Figure(go.Treemap(
        ids=treemap_df['id'].tolist(),
        labels=treemap_df['labels'].tolist(),
        parents=treemap_df['parent'].tolist(),
        values=treemap_df['Sample points'].tolist(),
        branchvalues='total',
        text='R<sup>2</sup>: ' + filtered_df['R^2'].astype(str) + '<br>' + filtered_df['Details'],
        hovertext=filtered_df['Study'] + '<br>R<sup>2</sup>: ' + filtered_df['R^2'].astype(str) +
            '<br>Number of samples: ' + filtered_df['Sample points'].astype(str),
        hoverinfo='text',
        textfont=dict(
            size=15,
        ),
        marker=dict(
            colors=filtered_df['R^2'],
            colorscale='Viridis',
            colorbar=dict(title='R<sup>2</sup>'),
            showscale=True
        )
    )
)

fig4 = fig4.update_layout(
    autosize=False,
    width=900,
    height=600,
    margin=dict(
        l=0,
        r=0,
        b=30,
        t=60,
    ),
    title='Figure S4: R<sup>2</sup> values across studies'
)

fig4.show()

# %% [code]
filtered_df['Variance'] = (4*filtered_df['R^2'])*((1-filtered_df['R^2'])**2)/filtered_df['Sample points']

metafor = importr('metafor')
stats = importr('stats')

metastudy = {}
for m in filtered_df.Measure.unique():
    nstudies=len(filtered_df.Measure[filtered_df.Measure==m])
    if nstudies > 2:
        df_m = filtered_df[filtered_df.Measure==m]
        df_m = df_m.sort_values(by=['Year'])
        
        r2 = rpy2.robjects.FloatVector(df_m['R^2'])
        var = rpy2.robjects.FloatVector(df_m['Variance'])
        fit = metafor.rma(r2, var, method="REML", test="knha")
        res = stats.predict(fit)
        
        results = dict(zip(res.names,list(res)))
        
        metastudy[m] = dict(pred=results['pred'][0], cilb=results['pred'][0]-results['ci.lb'][0],
                            ciub=results['ci.ub'][0]-results['pred'][0],
                            crub=results['cr.ub'][0], 
                            crlb=results['cr.lb'][0])

# %% [code]
measure_type_reverse={m:t for t,mlist in measure_type.items() for m in mlist}
fig5 = make_subplots(rows=3, cols=3, start_cell="top-left", vertical_spacing=0.05,
                      horizontal_spacing=0.2, x_title='R<sup>2</sup>',
                      subplot_titles=sorted(metastudy.keys(), key=measure_type_reverse.get))

row=1
col=1
for m in sorted(metastudy.keys(), key=measure_type_reverse.get):
    fig5.add_trace(go.Scatter(
        x=[round(metastudy[m]['crlb'],2) if round(metastudy[m]['crlb'],2)>0 else 0,
           round(metastudy[m]['crub'],2) if round(metastudy[m]['crub'],2)<1 else 1],
        y=['Mixed model','Mixed model'],
        line=dict(color='black', width=2, dash='dot'),
        hovertemplate = 'Prediction boundary: %{x}<extra></extra>',
        marker_symbol = 'hourglass-open', marker_size = 8
    ), row=row, col=col)
    
    fig5.add_trace(go.Scatter(
        x=[round(metastudy[m]['pred'],2)],
        y=['Mixed model'],
        mode='markers',
        marker = dict(color = 'black'),
        marker_symbol = 'diamond-wide',
        marker_size = 10,
        hovertemplate = 'R<sup>2</sup> estimate: %{x}<extra></extra>',
        error_x=dict(
            type='data',
            arrayminus=[round(metastudy[m]['cilb'],2) if round(metastudy[m]['cilb'],2)>0 else 0],
            array=[round(metastudy[m]['ciub'],2) if round(metastudy[m]['ciub'],2)<1 else 1])
    ), row=row, col=col)
    
    df_m = filtered_df[filtered_df.Measure==m]
    df_m = df_m.sort_values(by=['Year'], ascending=False)
    fig5.add_trace(go.Scatter(
        x=df_m['R^2'],
        y=df_m['Study'],
        text=df_m['Sample points'],
        customdata=df_m['Histology/microscopy measure'],
        mode='markers',
        marker = dict(color = color_dict[measure_type_reverse[m]]),
        marker_symbol = 'square',
        marker_size = np.log(50/df_m['Variance']),
        hovertemplate = '%{y}<br>R<sup>2</sup>: %{x}<br>Number of samples: %{text}<br>' +
            'Reference: %{customdata}<extra></extra>',
        error_x=dict(
            type='data',
            array=2*np.sqrt(df_m['Variance']))
    ), row=row, col=col)

    if col == 3:
        col = 1
        row += 1
    else:
        col += 1

fig5.update_xaxes(range=[0, 1])

fig5.update_layout(showlegend=False,
    title='Figure S5: Forest plots and mixed modelling results',
    width=1000,
    height=1400)
fig5.show()

# %% [code]
multcomp = importr('multcomp')
base = importr('base')

thres = filtered_df.Measure.value_counts() > 2
df_thres = filtered_df[filtered_df.Measure.isin(filtered_df.Measure.value_counts()[thres].index)]
r2 = rpy2.robjects.FloatVector(df_thres['R^2'])
var = rpy2.robjects.FloatVector(df_thres['Variance'])
measure_v = rpy2.robjects.StrVector(df_thres['Measure'])
measure_f = rpy2.robjects.Formula('~ -1 + measure')
env = measure_f.environment
env['measure'] = measure_v
study_v = rpy2.robjects.StrVector(df_thres['Study'])
study_f = rpy2.robjects.Formula('~ 1 | study')
env = study_f.environment
env['study'] = study_v
fit_mv = metafor.rma_mv(r2, var, method="REML", mods=measure_f, random=study_f)

glht = multcomp.glht(fit_mv, base.cbind(multcomp.contrMat(base.rep(1,9), type="Tukey")))
mtests = multcomp.summary_glht(glht, test=multcomp.adjusted("bonferroni"))
mtest_res = dict(zip(mtests.names, list(mtests)))
stat_res = dict(zip(mtest_res['test'].names, list(mtest_res['test'])))
pvals = stat_res['pvalues']
zvals = stat_res['tstat']

# %% [code]
measure_list = df_thres['Measure'].unique()
measure_list.sort()
n = len(measure_list)

pvals_list = []
zvals_list = []
idx = 0
for i in range(n):
    pvals_i = [0] * i
    pvals_i.append(np.nan)
    pvals_i.extend(pvals[idx:idx+n-i-1])
    zvals_i = [0] * i
    zvals_i.append(np.nan)
    zvals_i.extend(zvals[idx:idx+n-i-1])
    idx = idx + n - i - 1
    pvals_list.append(pvals_i)
    zvals_list.append(zvals_i)

pvals_list=np.array(pvals_list)    
zvals_list=np.array(zvals_list)
pvals_list=pvals_list+pvals_list.T
zvals_list=zvals_list-zvals_list.T
    
fig6 = make_subplots(rows=1, cols=2, horizontal_spacing=0.2,
                     subplot_titles=['z-scores',
                                     'p-values'
                                    ])

fig6.add_trace(go.Heatmap(
    z=zvals_list,
    x=measure_list,
    y=measure_list,
    customdata=pvals_list,
    hovertemplate = '%{x}-%{y}<br>z-score: %{z:.2f}<br>p-value: %{customdata:.5f}<extra></extra>',
    hoverongaps = False,
    colorscale='RdBu',
    colorbar=dict(title='z-score', x=0.42),
    showscale=True
), col=1, row=1)

fig6.add_trace(go.Heatmap(
    z=pvals_list,
    x=measure_list,
    y=measure_list,
    customdata=zvals_list,
    hovertemplate = '%{x}-%{y}<br>z-score: %{customdata:.2f}<br>p-value: %{z:.5f}<extra></extra>',
    hoverongaps = False,
    colorscale='Purples',
    colorbar=dict(title='p-value'),
    zmin=0,
    zmax=0.05,
    showscale=True,
    reversescale=True
), col=2, row=1)

fig6.update_layout(
    title='Figure S6: Statistical pairwise comparisons between R<sup>2</sup> estimates',
    height=500,
    width=950,
    paper_bgcolor='rgba(0,0,0,0)',
    plot_bgcolor='rgba(0,0,0,0)',
)

fig6.show()

# %% [code]
structures={'Lesions':'Lesions',
            'Substantia nigra':'Deep grey matter',
            'Hippocampal commissure':'White matter',
            'Putamen':'Deep grey matter',
            'Motor cortex':'Grey matter',
            'Globus pallidus':'Deep grey matter',
            'Perforant pathway':'White matter',
            'Mammilothalamic tract':'White matter',
            'External capsule':'White matter',
            'Inter-peduncular nuclues':'Deep grey matter',
            'Hippocampus':'Deep grey matter',
            'Thalamic nuclei':'Deep grey matter',
            'Thalamus':'Deep grey matter',
            'Cerebellum':'Grey matter',
            'Amygdala':'Deep grey matter',
            'Cingulum':'White matter',
            'Striatum':'Deep grey matter',
            'Accumbens':'Deep grey matter',
            'Basal ganglia':'Deep grey matter',
            'Anterior commissure':'White matter',
            'Cortex':'Grey matter',
            'Fimbria':'White matter',
            'Somatosensory cortex':'Grey matter',
            'Dorsal tegmental tract':'White matter',
            'Superior colliculus':'Deep grey matter',
            'Fasciculus retroflexus':'White matter',
            'Optic nerve':'White matter',
            'Dentate gyrus':'Grey matter',
            'Corpus callosum':'White matter',
            'Fornix':'White matter',
            'White matter':'White matter',
            'Grey matter':'Grey matter',
            'Optic tract':'White matter',
            'Internal capsule':'White matter',
            'Stria medullaris':'White matter'}


tissue_types=[]
for s in filtered_df['Specific structure(s)']:
    t_list=[]
    for i in s.split(','):
        t_list.append(structures[i.strip()])
    tissue_types.append('+'.join(list(set(t_list))))

filtered_df['Tissue types']=tissue_types

# %% [code]
fig7 = make_subplots(rows=3, cols=1, start_cell="top-left", vertical_spacing=0.2, y_title='R<sup>2</sup>',
                     subplot_titles=['R<sup>2</sup> values and reference techniques',
                                     'R<sup>2</sup> values and pathology',
                                     'R<sup>2</sup> values and tissue types'])

references = ['Histology', 'Immunohistochemistry', 'Microscopy', 'EM']

for r in references:
    df_r=filtered_df[filtered_df['Histology/microscopy measure'].str.contains(r)]
    fig7.add_trace(go.Box(
        y=df_r['R^2'],
        x=df_r['Histology/microscopy measure'],
        boxpoints='all',
        text=df_r['Measure'] + ' - ' + df_r['Study'],
        name=r
    ), col=1, row=1)

for t in filtered_df['Condition'].unique():
    df_t=filtered_df[filtered_df['Condition']==t]
    fig7.add_trace(go.Box(
        y=df_t['R^2'],
        x=df_t['Condition'],
        boxpoints='all',
        text=df_t['Measure'] + ' - ' + df_t['Study'],
        name=t
    ), col=1, row=2)

for t in filtered_df['Tissue types'].unique():
    df_t=filtered_df[filtered_df['Tissue types']==t]
    fig7.add_trace(go.Box(
        y=df_t['R^2'],
        x=df_t['Tissue types'],
        boxpoints='all',
        text=df_t['Measure'] + ' - ' + df_t['Study'],
        name=t
    ), col=1, row=3)   

fig7.update_layout(
    title='Figure S7: Experimental conditions and methodological choices influencing the R<sup>2</sup> values',
    showlegend=False,
    height=1200,
    width=900
)

fig7.show()

# %% [code]
fig8 = make_subplots(rows=2, cols=2, start_cell="top-left", vertical_spacing=0.15, y_title='R<sup>2</sup>',
                     subplot_titles=['R<sup>2</sup> values and magnetic fields',
                                     'R<sup>2</sup> values and tissue conditions',
                                     'R<sup>2</sup> values and co-registration',
                                     'R<sup>2</sup> values and human/animal tissue'
                                    ])

for m in measure_type.keys():
    df_m=filtered_df[filtered_df["Measure"].isin(measure_type[m])]
    fig8.add_trace(go.Scatter(x=df_m['Magnetic field'],
                              y=df_m['R^2'],
                              text=df_m['Measure'] + ' - ' + df_m['Study'],
                              marker=dict(color=color_dict[m]),
                              name=m,
                              mode='markers'), col=1, row=1)

fig8.update_layout(
    xaxis=dict(title='Magnetic field [T]')
)

for t in filtered_df['Tissue condition'].unique():
    df_t=filtered_df[filtered_df['Tissue condition']==t]
    fig8.add_trace(go.Box(
        y=df_t['R^2'],
        x=df_t['Tissue condition'],
        boxpoints='all',
        text=df_t['Measure'] + ' - ' + df_t['Study'],
        name=t
    ), col=2, row=1)
    
for t in filtered_df['Co-registration'].unique():
    df_t=filtered_df[filtered_df['Co-registration']==t]
    fig8.add_trace(go.Box(
        y=df_t['R^2'],
        x=df_t['Co-registration'],
        boxpoints='all',
        text=df_t['Measure'] + ' - ' + df_t['Study'],
        name=t
    ), col=1, row=2)
    
for t in filtered_df['Human/animal'].unique():
    df_t=filtered_df[filtered_df['Human/animal']==t]
    fig8.add_trace(go.Box(
        y=df_t['R^2'],
        x=df_t['Human/animal'],
        boxpoints='all',
        text=df_t['Measure'] + ' - ' + df_t['Study'],
        name=t
    ), col=2, row=2)

fig8.update_layout(
    title=dict(text='Figure S8: Other factors to consider when assessing R<sup>2</sup>', x=0.1),
    margin=dict(l=100),
    showlegend=False,
    height=900,
    width=900
)    
    
fig8.show()
```
